# Effects of temperature on the life-history traits of *Myzus persicae* and its efficiency in transmitting potato virus Y (PVY) in potato crops

**DOI:** 10.1101/2024.10.05.616836

**Authors:** Bonoukpoè M. Sokame, Henri E.Z. Tonnang, Heidy Gamarra, Pablo Carhuapoma, Jan Kreuze, Ali Arab, Peter A. Armbruster, Leah R. Johnson, Oswaldo C. Villena

## Abstract

Aphids are highly sensitive to temperature changes and play a crucial role in transmitting plant viruses, accounting for the transmission of more than 50% of viruses that cause disease in crops. Among them, *Myzus persicae* is a major global pest, affecting over 400 plant species and transmitting more than 100 plant viruses, including potato virus Y (PVY), which poses a severe threat to potato crops. This study examines how temperature influences the life-history traits of *M. persicae* and its efficiency in transmitting PVY. Our research revealed that temperature significantly affects developmental duration, survival, and fecundity of *M. persicae*. The aphids exhibit the longest lifespan at 10°C and the shortest at 30°C. Similarly, fecundity declined from 29.81 offspring per female at 10°C to 14.25 at 30°C. PVY transmission efficiency was highest at 20°C. We also mapped potential PVY transmission regions and identified tropical and subtropical areas as high-risk due to their favourable temperatures and aphid’s abundance. Understanding these geographical variations is crucial for effective pest/disease management. Our findings emphasize the importance of integrating climatological, ecological, and epidemiological data to develop robust pest/disease management strategies to mitigate the impact of *M. persicae* and PVY on potato production and enhancing global food security.

## 1. Introduction

The green peach aphid, *Myzus persicae* Sulzer (Hemiptera: Aphididae), is a globally significant pest that affects a wide range of crops due to its ability to feed on over 400 plant species and transmit more than 100 plant viruses [1]. Among the crops it severely impacts, potatoes are particularly vulnerable due to the aphid’s role in transmitting various viruses amongst which the most important is Potato virus Y (PVY), a pathogen that significantly reduces yield and quality. Understanding the dynamics of *M. persicae* infestations and PVY transmission is critical for developing effective management strategies to protect potato production. *Myzus persicae* causes direct damage to potato plants through its feeding activity. Aphids feed by inserting their stylets into the phloem of the plant and extracting sap, which weakens the plant, and can lead to symptoms such as chlorosis, leaf curling, and reduced growth rates [2]. Heavy infestations can cause significant reductions in photosynthesis and overall plant vigour, ultimately impacting tuber yield and quality. Yield losses of up to 50% in potato crops and increased deformities and reduced size, with quality losses up to 40%, have been reported [3]. However, the more insidious damage comes from the transmission of plant viruses such as PVY. *Myzus persicae* is an efficient vector of PVY, spreading the virus as it feeds on potato plants [2]. PVY is one of the most economically damaging viruses affecting potatoes. It belongs to the *Potyviridae* family and is responsible for causing a range of symptoms including mottling, leaf drop, and tuber necrosis, which significantly diminish the marketability and yield of potato crops [4]. Infected plants often produce fewer and smaller tubers, and tubers may have necrotic rings, making them unmarketable. Estimates suggest that PVY can reduce potato yields by up to 80% in severely affected fields [5]. The economic losses are compounded by the increased costs associated with managing the disease and the reduced marketability of infected potatoes.

To mitigate the impact of *M. persicae* and PVY, integrated pest management (IPM) strategies are essential. These strategies include the use of certified virus-free seed potatoes, crop rotation, and the timely application of insecticides to control aphid populations [6]. Biological control agents, such as natural predators and parasitoids, also reduce aphid numbers. Additionally, the deployment of PVY-resistant potato varieties is effective in reducing the incidence of the disease [4]. Despite advances in IPM, challenges remain in managing *M. persicae* and PVY due to factors such as insecticide resistance, the emergence of new PVY strains, the rapid non-persistent transmission mode of PVY, and, most importantly, climatic variability. Therefore, understanding the ecological and epidemiological interactions between aphids and PVY under changing environmental conditions remains a high priority. The study of *M. persicae*’s life history traits and its efficiency in transmitting PVY under varying temperature conditions is crucial for developing effective pest management strategies.

Temperature is a critical environmental factor that influences the biology, behaviour, and ecology of insects. For *M. persicae*, temperature affects developmental rates, survival, fecundity, and virus transmission efficiency [7]. Understanding these temperature-dependent life history traits is essential for predicting aphid population dynamics and virus spread under changing climatic conditions, which in turn can inform integrated pest management (IPM) practices [8,9]. Numerous studies have explored the influence of temperature on aphid biology. For instance, Khurshid et al. (2022) demonstrated that the developmental rates of *M. persicae* increase with temperature up to an optimal point, beyond which higher temperatures become detrimental to survival and reproduction [10]. This aligns with the findings of Baral et al. (2022), who demonstrated a similar trend in the development rates of *M. persicae* with increasing temperature [11]. Additionally, Jeger et al. (2004) reported that temperature variations significantly impact the transmission efficiency of plant viruses by aphids, underscoring the need for temperature-specific management strategies in pest control [12]. However, there is a need for comprehensive studies that integrate the effects of temperature on both the life history traits of *M. persicae* and its ability to transmit PVY. Such integrated assessments are vital for developing robust predictive models and effective management practices, particularly in the context of global climate change [13].

This study aims to address this gap by systematically investigating the influence of a range of temperatures on the survival, development, fecundity, and PVY transmission rates of *M. persicae*. We hypothesize that temperature significantly affects these parameters and that there is an optimal temperature range where *M. persicae* exhibits peak performance in terms of survival and reproduction, while extreme temperatures (both high and low) adversely affect these traits. Additionally, we anticipate that the efficiency of PVY transmission by *M. persicae* will exhibit a temperature-dependent pattern, potentially complicating disease management under fluctuating climatic conditions.

## 2. Materials and Methods

### 2.1 *Myzus percicae* life-history traits

#### a) Plant material and aphid colonies

For generating *M. persicae* life table data, a colony of *M. persicae* originating from potato crops in Kenya and maintained on potato plants for five years in a greenhouse facility at the International Centre of Insect Physiology and Ecology (*icipe*) was used for the experiments. The rearing conditions were controlled at 20−23 °C, 70–95% RH, and a 12:12 h light-dark photoperiod. The potato variety “Shangi,” known for its popularity and moderately high yields in Kenya, served as the host plant. Leaves from pesticide-free plants were used throughout the experiment. Before use, the leaves were disinfected in a 1% sodium hypochlorite solution for 5 minutes, followed by thorough rinsing with tap and distilled water [14,15]. This procedure ensured high-quality leaves for the aphids. During the experiments, leaf discs showing signs of chlorosis or dehydration were replaced, and the insects were carefully transferred to fresh discs using a paintbrush.

#### b) Development and survival

Individual females of aphid *M. persicae* were transferred to separate Petri dishes (10 cm diameter) containing a host plant leaf disc and 1% agar solution. The dishes were kept at 22 ± 1°C and 70 ± 10% relative humidity for 6 h; after that the females and all the nymphs, except for ten per dish, were removed. The dishes with newly hatched neonates were then incubated in environmental chambers (phytotron, Weiss Technik, Germany) at different constant temperatures 10, 15, 20, 25, and 30°C at 70 ± 10% relative humidity and a 12-hour photoperiod. Juvenile development and survival were monitored daily until they developed into adult aphids, the females of which were then used in the fertility study described below [14,15]. Ten replicates of a cohort of 10 nymphs per Petri dish were tested for each temperature.

#### c) Adult longevity and fecundity

A simple random sampling design was used, which included five temperature variables (10, 15, 20, 25, and 30 °C) and 100 repetitions. Female adult aphids were incubated at the appropriate temperature in dishes containing potato host plant leaf discs, maintained under a 12-hour photoperiod, and were transferred to new leaf discs when necessary [15]. The pre-reproductive and reproductive periods were evaluated under a stereomicroscope every 24 hours, and the number of nymphs produced, and their longevity was determined at each temperature. All nymphs produced were removed to eliminate the potential effect of crowding.

#### d) Key Population Parameters and Formulas

There are a variety of demographic summaries that capture multiple important aspects of population growth. Many of these are related to each other and are usually calculated together. Here we summarize the population metrics calculated in this study following Gao et al., 2012 [14]. The first four are the key life-history/demographic parameters on which the other metrics are built:

1) Age-stage specific survival rate (*Sxj*) represents the probability that an individual will survive to age *x* and stage *j*.

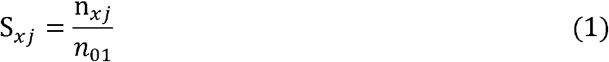

where *n*_*xj*_ is the number of individuals surviving to age x and stage j, and *n*_01_ is the initial number of neonates.
2) Age-specific survival rate *(lx)* indicates the probability that a newly laid neonate will survive to age x.

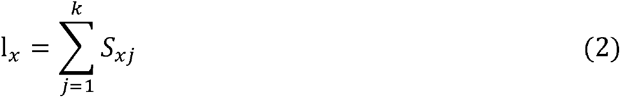

where *k* is the number of developmental stages.
3) Age-specific fecundity *(mx)* represents the mean number of offspring produced per individual at age x.

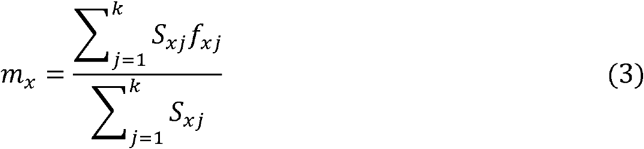

where *f*_*xj*_ is the fecundity of individuals at age x and stage j.
4) Age-stage life expectancy *(e*_*xj*_) indicates the expected lifespan of individuals at age *x* and stage *j*.

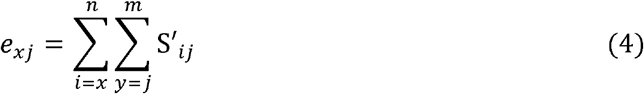

where *s*′_*ij*_ is the probability that an individual of age *x* and stage *j* will survive to age *i* and stage *y*. There are multiple summaries of population growth that can be built from the four demographic parameters above. Because these capture different aspects of population performance we choose to include the following five:

1) Net reproductive rate *(R*_*0*_) estimates the total number of offspring that an individual is expected to produce over its lifetime and is calculated as:

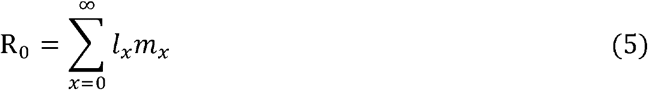
2) Intrinsic rate of increase *(r*_*m*_) is calculated using the Euler-Lotka equation, representing the growth rate of the population.

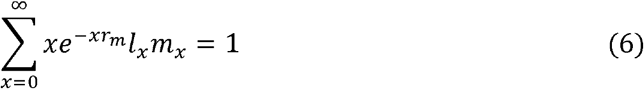
3) Finite rate of increase *(λ)* indicates the population’s growth rate per unit time.

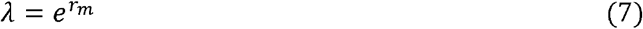
4) Mean generation time *(T)* represents the average time between the birth of an individual and the birth of its offspring and is calculated as follow:

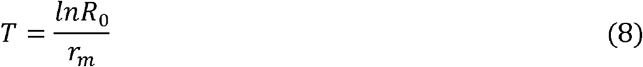
5) Doubling time *(DT)* is a key demographic parameter used to describe the period required for a population to double in size. The concept of doubling time is based on exponential growth, where the population increases at a constant rate over time.

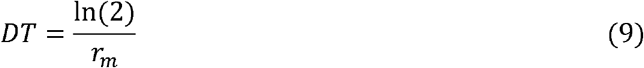

#### e) Data analysis

Count data (nymph longevity, adult longevity, total longevity, and fecundity expressed as offspring per female) and life table parameters data (Net reproduction rate (*R*_*o*_), Intrinsic rate of increase (*r*_*m*_), Finite rate of increase (*λ*), Mean generation time (*T*), Doubling time (*DT*)) were tested for normality using Shapiro-Wilk test and homogeneity of variance using Levene test. The data were not normally distributed, and variances were not homogeneous. Count data were therefore, analysed with generalized linear model (GLM) with negative binomial error distribution considering overdispersion. Whenever there was a significant difference, the means were separated using Tukey’s honest significant difference (HSD) test using “*agricolae*” package in R [16,17]. Life table parameter data were analysed using Kruskal-Wallis nonparametric procedure. The survival and life expectancy curves for immature life stage and adult of *M. percicae* were generated using Kaplan-Meier estimator method, and log-rank test was used to compare the effect of temperature on *M. persicae* immature and adult in SigmaPlot software version 14.0 (Systat Software, Inc., San Jose, California).

### 2.2. Myzus persicae phenology model development and analysis

The development of the *M. persicae* phenology model simulation was conducted using the Insect Life Cycle Modelling (<u>ILCYM</u>) software version 4.0, developed by CIP [18]. ILCYM is a software that facilitates the development of pest insect phenology models and provides analytical tools for studying pest population ecology. The ILCYM software, which employs R statistics [17] for all calculations, is freely available online. Development time, development rate, senescence, mortality, and total oviposition data collected under different constant temperature conditions were included in the phenology model. The best-fit model was selected based on the Akaike Information Criterion (AIC), with lower AIC values indicating better fit [19,20]. Data were transformed into interval censored time-to-event data for survival analysis. Parametric accelerated failure time (AFT) modelling was used to determine medians and the distribution of development times, with models adjusted using the survreg procedure in R [21,22].

For development times and adult longevity, log-error distributions were assumed, and the most appropriate distribution link function (log-logistic, lognormal, or Weibull) was chosen based on maximum likelihood. These functions were fitted in terms of ln-times. Lower developmental thresholds and thermal requirements were calculated using linear regression between temperature and observed development rates within the linear range. Survival time of immature stages was calculated from the relative frequency of surviving insects, and different nonlinear models were adjusted by regression to describe mortality rates and fecundity by temperature. Development rate was expressed as the reciprocal of mean development times, and mortality was calculated from cohort mortality frequency.

### 2.3 Transmission rates of potato virus Y (PVY) by *Myzus persicae*

#### a) Plant material, aphid colony, and PVY strain

For the virus transmission experiments, a colony of *M. persicae* was isolated and maintained on potato plants for six years in a rearing facility (temperature-controlled greenhouse) at the International Potato Center (CIP) in La Molina, Lima, Peru, at a temperature between 20-23°C, 70-95 % RH, and a photoperiod of 12:12 hours light(L): dark(D). Virus-free potato plants were obtained from the CIP’s germplasm bank in Lima, Peru (accession CIP720201; cultivar “Yungay”). PVY virus was sourced from the CIP potato virus collection, propagated, and maintained in potato plants. Infected tubers served as source plants for the transmission studies, with virus infection confirmed approximately 25 days post-planting by RT-PCR. Healthy sprouted potatoes, prepared as targets for inoculation experiments, were planted 7 days before testing and maintained under the same conditions as the virus transmission experiments.

#### b) Virus transmission experiments

A diagram summarising the virus transmission experiments is provided in Figure S2. PVY transmission by *M. persicae* was determined at 4 constant temperatures of 12°,15°, 20°, and 25 °C (± 0.5 °C). For each temperature, 10 single aphid transmissions (described below) were performed per replication and replicated 3 times per temperature initially. Aphid transmissions were performed with a fasting period where the insect does not feed on the plant, an acquisition access period (AAP) to acquire the virus from the infected plants and inoculation access period (IAP) to transmit acquired virus to target plants (Figure S2). A 2-hour fasting period was required before starting the acquisition access period. During the AAP, PVY-infected potato plants (1 plant per 30 insects) were placed inside an acrylic box, covered with nylon mesh, in a growth chamber under each of the 4 predetermined temperatures. Adult aphids were caged with a clip-cage (17 × 13 × 13 cm) covered with nylon mesh to three leaves per plant (10 adults/leaf), one from the apical part and two from the middle part of the plant and were allowed to feed on the plants during this 5min AAP. After this period, each of these adults were transferred individually directly to a single 10-day old healthy plant for a 24-h IAP covered by nylon mesh in growth chambers at the same temperature as during the AAP. Each batch of 10 insects that acquired the virus from one leaf was considered a replication.

After the 24-hour IAP, the insects were removed from the cages and the plants treated with the insecticide Spirotetramat at 0.1 % to ensure that no insects remained. The inoculated plants were subsequently maintained in a greenhouse at a photoperiod of 12:12 hours L: D and temperature of 18°C during the light and dark period, for 25 days. After this 25-day incubation period, virus presence was assessed taking systemic leaves (non-inoculated leaves) submitted to RT-PCR analysis.

#### c) Reverse transcription polymerase chain reaction (RT-PCR)

Total RNA was extracted from sampled leaves using the CTAB method (Adapted from Lodhi et al 1994). The cDNA for RT-PCR was synthesized as follows: 10 μl of nuclease-free water (NFW), 1 μl of Random primers (250 ng/μl) and 1 μl of total RNA (500 ng/μl) were mixed and denatured in a thermocycler (Applied Biosystems, Foster City, City, CA) at 65 °C for 10 min and then cooled to 10 °C for 5 min. Subsequently, RT Mix (4 μl 5X First Strand Buffer, 0.5 μl dNTPs 10 mM, 0.5 μl DTT 100 mM, 0.5 μl RNAse OUT and 0.5 μl M-MLV reversed transcriptase), was added and then incubated at 37 °C for 50 min, followed by 95 °C for 15 min and finally cooled to 10 °C. The final volume obtained was a 20 μl cDNA, which was diluted with 40 μl of NFW to obtain a dilution of 1:5 for subsequent PCR. The reaction conditions were as follows: 4 μL of PCR 5X Buffer (Promega), 1 μl of MgCl2 (25 mM), 0.5 μl of dNTPs (10 mM), 0.5 μL of Primers (PVY_F /PVY_R) both at 10 μM, 0.125 μl of the Taq polymerase (Promega), 8.375 μl of nuclease-free water (NFW) and 5 μl of diluted cDNA. The mixture was placed in a thermal cycler with the following conditions: one cycle at 95 °C for 2 min, followed by 35 cycles at 94 °C for 30 sec, 57 °C for 45 sec, and 72 °C for 45 sec followed by 10 min at 72 °C. Amplified bands were visualised in a 1% agarose gel stained with Gelred (Invitrogen) under UV light.

#### d) Developing the mathematical model for virus transmission efficiency

The transmission evaluated for each adult in a n_j_ sample at the j^th^ temperature *T*_*j*_ was considered a dichotomous variable y_ij_, with values of 1 for success and 0 for failure, for the i^th^ insect at the j^th^ temperature. This approach allowed for the calculation of the absolute frequency or total number of insects that transmitted the virus at each temperature in y_ij_, which give the percentage of transmission. This yielded a transmission rate for each constant temperature assessed, with temperature as the independent variable and percentage of transmission as the dependent variable. The relationship between these variables is represented by a nonlinear function *f(T) = p*. We tested approximately 20 models available in ILCYM, which were developed to describe insect transmission rates, and selected the best models based on the corrected Akaike Information Criterion (AICc). Parameters for each model were automatically estimated by ILCYM using the Levenberg-Marquardt algorithm [23], implemented through the “nls.lm” function (contained in the R package “minpack.lm”), which minimizes the sum of squares of the vectors returned for each function. Several models demonstrated a significant overall fit; finally, we used the Taylor model (Table 6). Therefore, all candidate models were implemented as optional functions within the ILCYM package for analysing temperature-dependent vector virus transmission probability (efficiency) in insect vectors [20,24,25]. All statistical calculations and model implementation coding were conducted in R-3.4.1 [17].

### 2.4. Risk Index Mapping

To implement and assess the risk of *M. persicae* using its phenological model in a GIS (Geographic Information System) environment, the potential distribution and risk mapping module of the ILCYM 4.0 software was used, which allows for spatial simulations at regional and global scales. As such, it is then possible to map vulnerability of geographic areas to the possible presence of this pest, estimating the potential for the insect to establish itself and it’s possible abundance, following the methodology described by Sporleder et al. (2017, 2023) and Kroschel et al. (2013, 2016) [17,23,26,27]. The risk of establishment is then estimated by calculating the Establishment Index (EI) and the potential abundance by calculating the Generation Index (GI), for regions where potatoes are produced at a global level.

### 2.5. Virus Transmission Index

ILCYM can calculate the virus transmission index (VTI) of the insect vector. These indices help identify areas vulnerable to pest infestation that have the capacity to transmit a certain virus, thus allowing the development of a risk alert system through the interpretation of the critical values produced by the software, following the methodology described by Gamarra et al. (2020) [28].

ILCYM calculates a specific transmission risk using the values obtained from immature survival, reproduction, transmission rate and their respective formulas as follows:

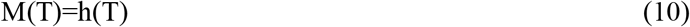

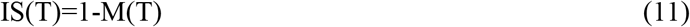

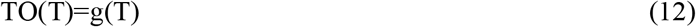

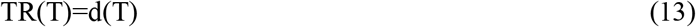

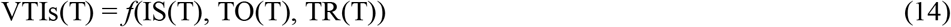

where, *T* is the air temperature, *M* is the mortality (Total mortality of all immature stages), *IS* is the immature survival, *TO* is the total oviposition, *TR* is the transmission rate.

Hence:

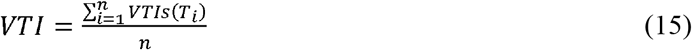

n=number of temperature records for 1 full year

To develop global level transmission risk maps, ILCYM simultaneously extracted the monthly maximum and minimum temperature data for one year (12 sets of monthly data starting from January to December) with their respective geographical coordinates from the WorldClim 2.1 database (https://worldclim.org/data/worldclim21.html).

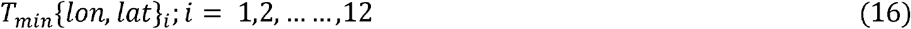

This represents the minimum temperature where, “*lon*” is the longitude and “*lat*” is the latitude respectively.

Temperature data were arranged into 24 matrices based on longitude and latitude, split into 12 for minimum and 12 for maximum temperatures. Each geographical point was then represented in a table with two columns, holding minimum and maximum temperatures for each month. These tables facilitated spatial phenological simulations. Correspondingly, for every risk index calculated, an additional matrix was generated with identical longitude and latitude organization. These matrices were then converted into ASCII files and imported into the R programming environment, which was equipped with GIS capabilities, for the purpose of mapping and visualize the VTI index.

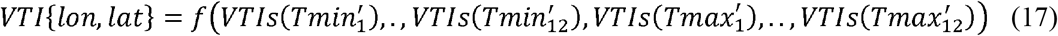

where, 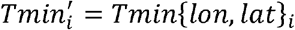and 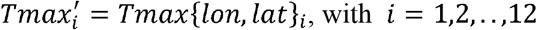

The distribution of *M. persicae* in potato-growing areas was assessed using the potato production areas for the world (CIP, 2010). Areas were mapped with a 10-minute grid resolution to identify the regions at risk of transmission based on the potential establishment of *M. persicae* populations, considering the presence of potatoes.

## 3. Results

### a) *Myzus persicae* nymphal longevity, survival, and development rate

A comprehensive analysis of the development time, longevity, and survival of the nymphal stage of the aphid *M. persicae* at temperatures ranging from 10°C to 30°C was conducted using GLM models (Table 1) and the ILCYM software (Table S1). For nymph longevity, the longest duration was observed at 10°C, averaging 12.58 days, which decreased steadily with increasing temperatures, reaching the lowest at 30°C with an average of 9.74 days. Nymph survival was lowest at extreme temperatures, with 58% survival at 10°C and 46% survival at 30°C (Table S1).

**Table 1.** Nymphal and adult developmental time and fecundity of aphid *Myzus persicae* under different temperatures.

| Temperature (°C) | Nymph longevity | Adult longevity | Total longevity | Fecundity (offspring per female) |
| --- | --- | --- | --- | --- |
| 10°C | 12.58 ± 0.38 **a** | 12.46 ± 0.53 **a** | 25.04 ± 0.56 **a** | 29.81 ± 1.83 **a** |
| 15°C | 11.50 ± 0.36 **b** | 13.48 ± 0.53 **a** | 24.48 ± 0.49 **b** | 45.82 ± 2.32 **b** |
| 20°C | 11.00 ± 0.26 **c** | 11.05 ± 0.50 **b** | 21.37 ± 0.56 **c** | 39.68 ± 2.08 **c** |
| 25°C | 10.32 ± 0.32 **d** | 9.21 ± 0.42 **c** | 18.96 ± 0.46 **d** | 38.59 ± 2.51 **c** |
| 30°C | 9.74 ± 0.33 **e** | 6.44 ± 0.40 **d** | 17.94 ± 0.54 **e** | 14.25 ± 1.09 **d** |
| LR$\chi^2$ | 26.11 | 111.98 | 104.81 | 124.29 |
| df | 4 | 4 | 4 | 4 |
| p | <0.0001 | p <0.0001 | p <0.0001 | p <0.0001 |

The variation in development times during the immature life stage across all temperatures was best described using the Weibull function (Table S1). This approach yielded highly significant common scale parameters (p < 0.001), as indicated by the lowest AIC values, effectively capturing the variability in nymph development with respect to temperature. The common slopes were also highly significant (p < 0.001), providing an adequate description of the overall variability in the development of immature life stages (Figure 1A). A linear model failed to describe the development rate appropriately at extreme temperatures, so a nonlinear model was fitted using AIC selection criteria. The temperature-dependent median developmental rate was accurately modelled by the Taylor model (Table S2). This model explained more than 97% of the variation in median development times due to temperature. The parameter *Topt* in the Taylor model indicated that the fastest development rate occurred at 23°C (Table S2, Figure 1B). Significant differences in mortality were observed across different temperatures (F = 3.04, df = 2, p = 0.247). Mortality rates were predicted to be highest at extreme temperatures, with 94% mortality at 0°C and 100% mortality at 45°C, while the lowest mortality was recorded at 20°C. The effects of temperature on the mortality of *M. persicae* in the immature stage was best described by a quadratic model (Table S3; Figure 1C). This model predicted increased mortality as temperatures deviated from the optimal range, indicating survival limits around 5°C and 35°C (Figure 1C).

**Figure 1.**
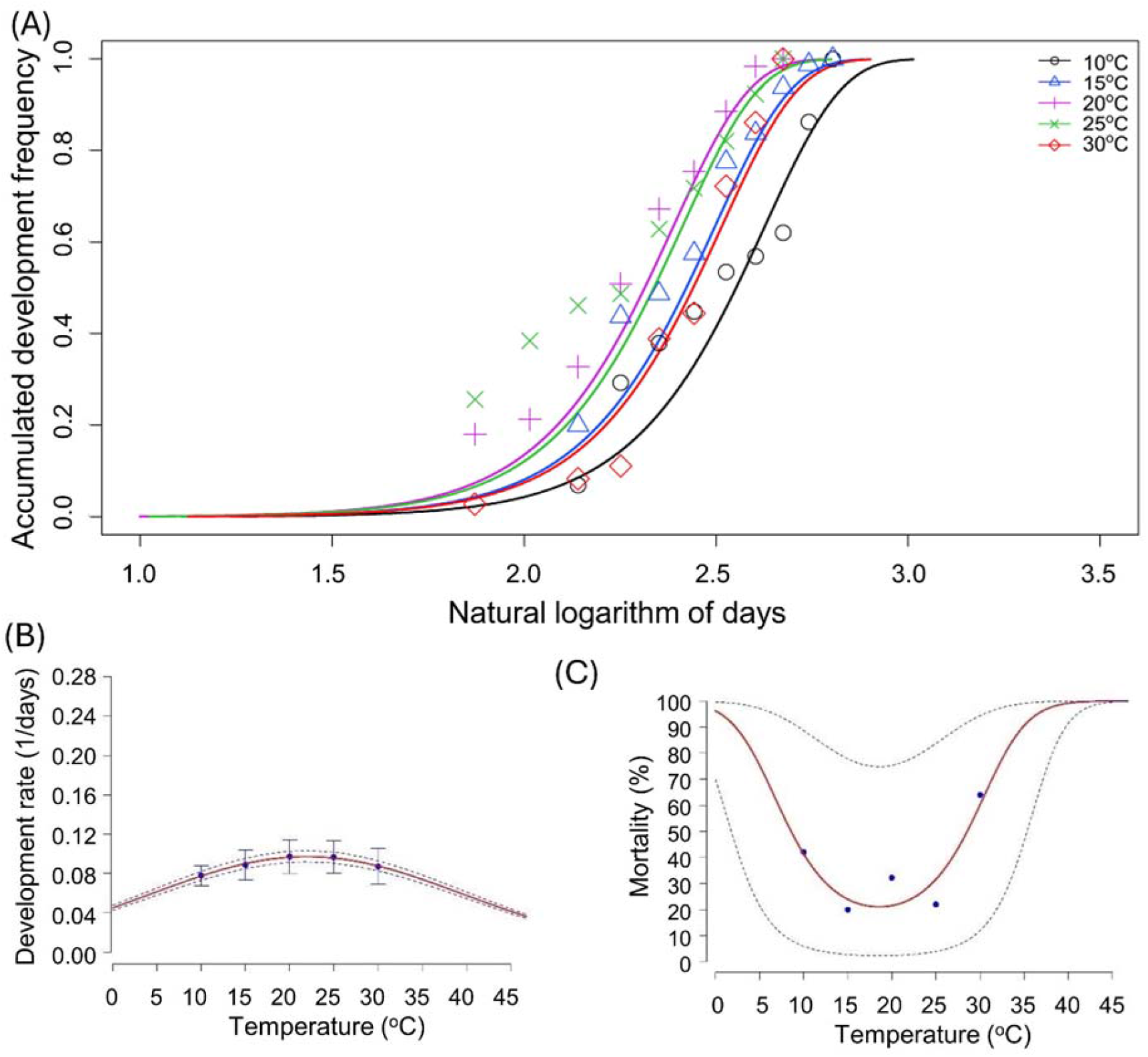
Temperature-dependent cumulative distribution of development time, development rate and mortality rate of nymph life stage of *Myzus persicae*

Furthermore, temperature significantly influences survival trends (Figure 2A). At 10°C, nymphs exhibit the highest resilience, maintaining nearly 100% survival until around day 9, after which their survival gradually declines to 0% by day 19 (Figure 2A). As the temperature increases, the survival rate declines more rapidly. At 15°C, the survival rate drops below 10% by day 15 and reaches 0% by day 26 (Figure 2A). This decline becomes sharper at 20°C, 25°C, and 30°C, with survival falling to 0% by day 14, 16, and 15, respectively, highlighting the critical temperature sensitivity of the nymph stage to increased thermal conditions (Figure 2A). Figure 2B presents life expectancy trends of *M. persicae* nymphs across temperatures ranging from 10°C to 30°C. Life expectancy decreases with age, but the rate of decline is influenced by temperature. At 10°C, life expectancy starts the highest, suggesting slower developmental rates or better survival at lower temperatures. As temperature increases, life expectancy starts lower and decreases more rapidly. At 30°C, nymphs show the lowest initial life expectancy, which rapidly declines, reaching near zero by about day 15 (Figure 2B).

**Figure 2.**
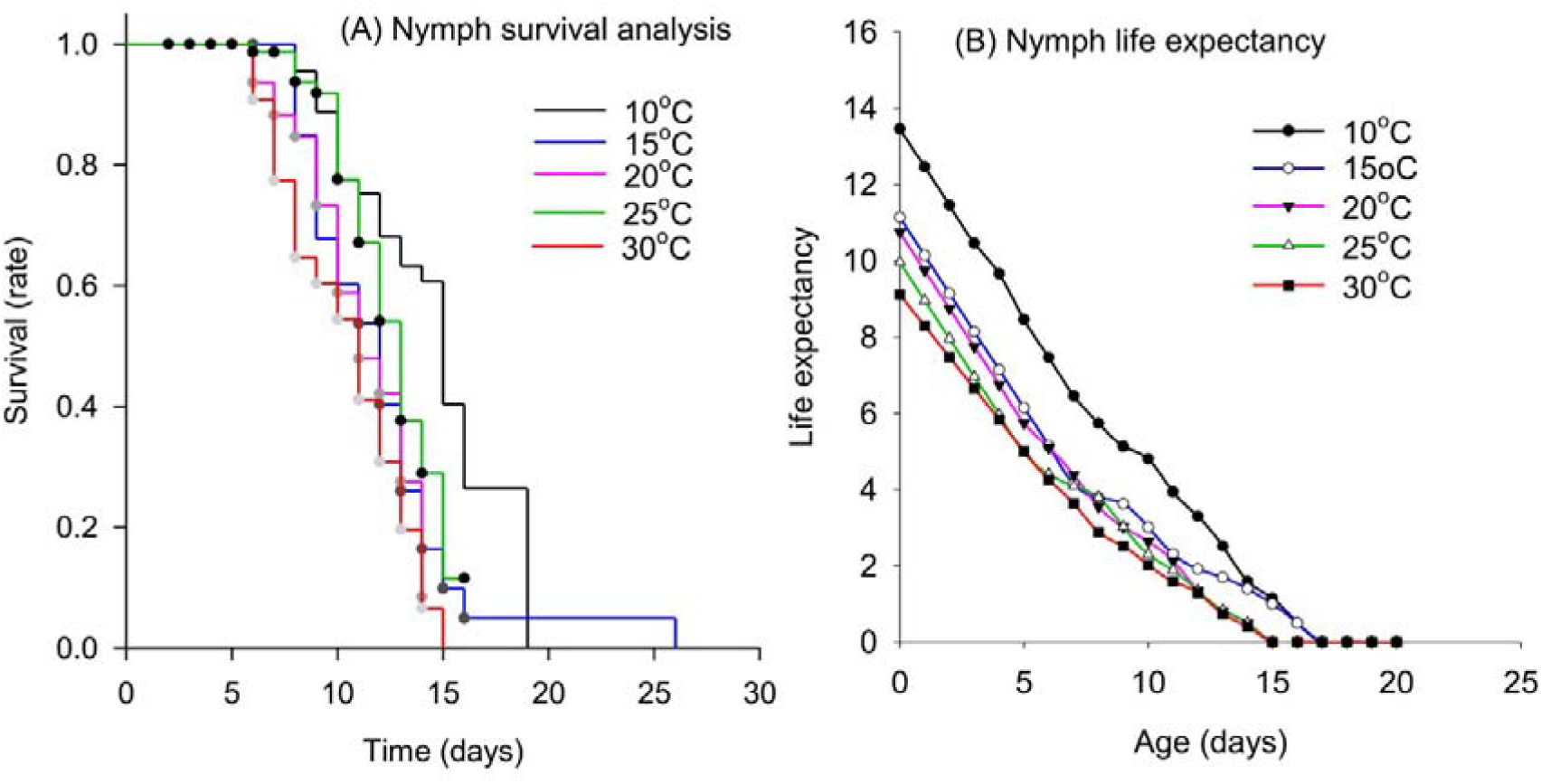
Temperature-dependent survival rate and life expectancy of nymph life stage of *Myzus persicae*

### b) *Myzus persicae* adult longevity, survival, and life expectancy

For adult longevity, the longest duration is at 15°C (13.48 days), which is not significantly different from 10°C, and the shortest at 30°C (6.44 days). Total longevity also decreases with temperature, from 25.04 days at 10°C to 17.94 days at 30°C (Table 1). Similar results and trends were obtained when data was analysed using ILCYM (Table S1).

Figure 3A indicates that the development time increased with temperature up to a certain point, beyond which it declined, where the temperature becomes detrimental to development; this is optimally described using a Weibull distribution function (Table S1). This approach produced highly significant common scale parameters (p < 0.001), as indicated by the lowest AIC values, effectively capturing the variability in adult development in relation to temperature. Additionally, the common slopes were highly significant (p < 0.001), offering a robust description of the overall variability in the development of the adult life stages. The survival rate was distinctly influenced by temperature, demonstrating clear temperature-dependent survival patterns (Figure 3B). At a lower temperature of 10°C, adults exhibit a gradual decline in survival, maintaining over 50% survival up to about day 12, and then decreasing to approximately 25% survival by day 16, and reaching 0% at day 22 (Figure 3B). As temperatures increase, a marked acceleration in the rate of decline in survival is observed. At 20°C, survival falls to about 50% at day 11 and reached 0% at day 18. Similarly, at 25°C, survival falls to about 50% at day 11 and falls below 20% at day 16 (Figure 3B). The trend becomes more pronounced at 30°C, where survival steepens further, dropping below 40% at day 7 and with survival plummeting to 0% by day 13 (Figure 3B). Similarly to immature life stage, life expectancy at adult stage at cooler temperatures (10°C and 15°C) starts higher and declines at a slower rate compared to warmer temperatures (Figure 3C). The life expectancy at 25°C and especially 30°C shows a steeper decline. At 30°C, adults have a significantly shorter life expectancy, with values plummeting to near zero by day 11 (Figure 3C).

**Figure 3.**
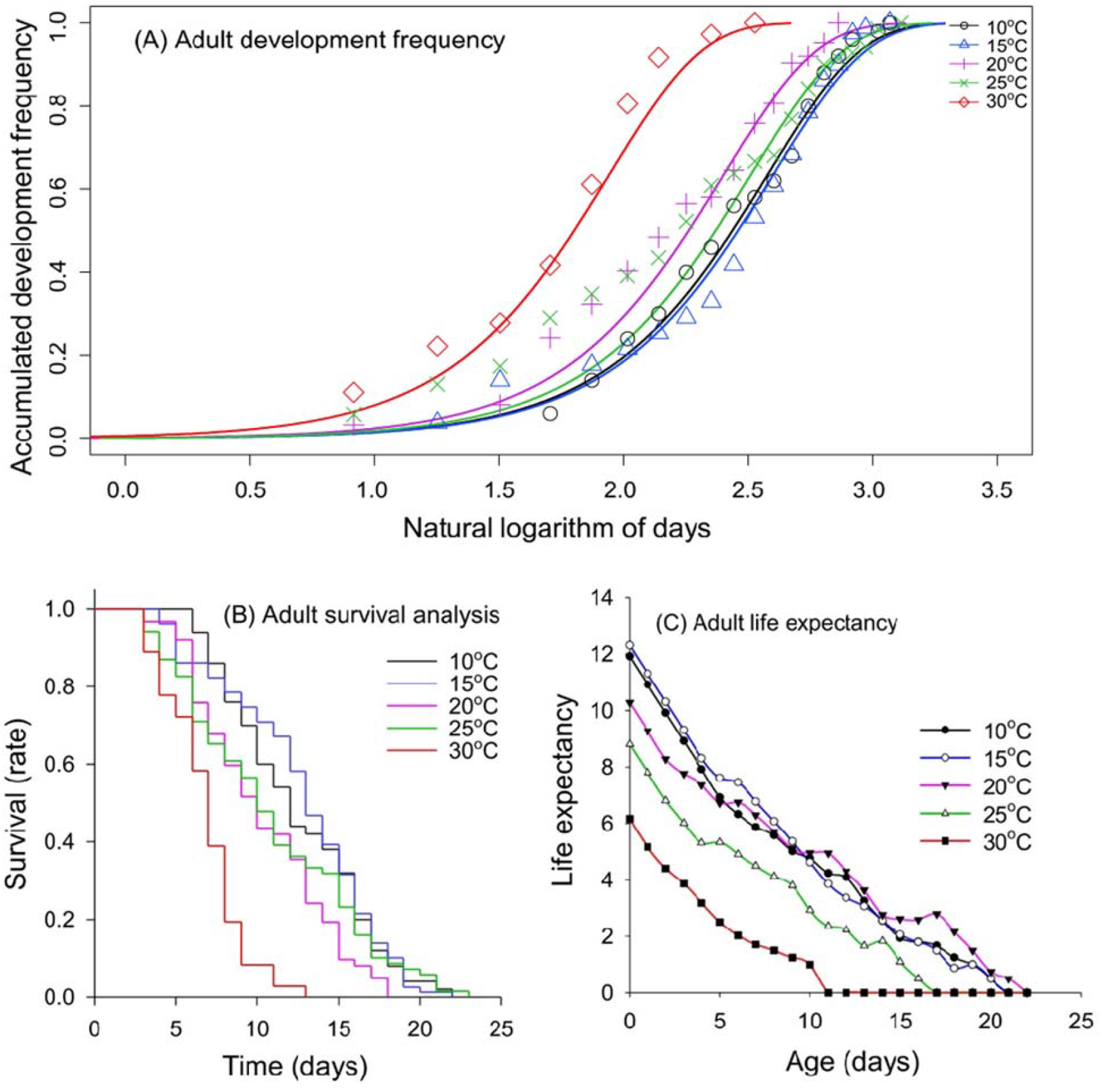
Temperature-dependent cumulative distribution of development time, survival rate and life expectancy of adult life stage of *Myzus persicae*

### d) Adult senescence rate and fecundity of *Myzus persicae*

A quadratic model was employed to explore the relationship between the senescence rates of adults and temperature (Figure 4A and Table S5). The lowest senescence rates were recorded within the temperature range of 10–20°C (Figure 4A). Female fecundity showed a significant decrease with increasing temperature either using a negative binomial GLM (Table 1) or the ILCYM software (Table S4). At 10°C, the average fecundity is 29.81 and 30.16 offspring per female respectively, which is the highest recorded, while at 30°C, it drastically drops to only 14.25 and 15.66 offspring per female respectively (Tables 1 and S4). This pattern suggests that lower temperatures favour higher reproductive output. The Taylor function demonstrated a significant impact of temperature on oviposition time. The effects of temperature on fecundity were best captured by a quadratic model, which predicted the highest fecundity at 15-20°C (Figure 4B and Table S5). Additionally, the relationship between temperature and both the survival time of adults and the oviposition rate were most accurately described by an exponential model (Figure 4C and Table S5).

**Figure 4.**
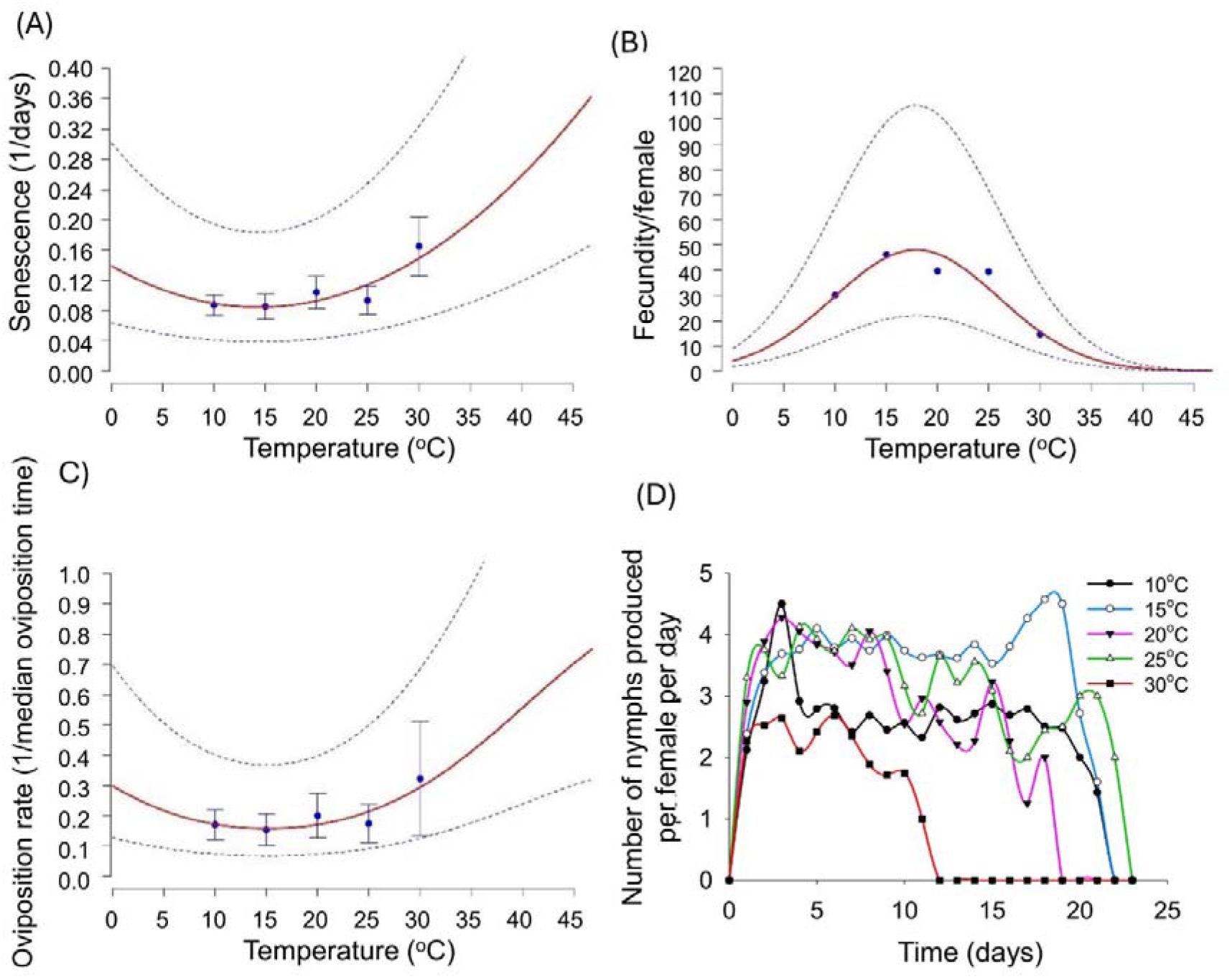
Temperature-dependent senescence rate, total fecundity per female, oviposition rate and daily fecundity of *Myzus persicae*

Figure 4D depicts the reproductive output, measured as the number of nymphs produced per female per day, of *M. persicae* across different temperature conditions. Each temperature condition, represented by distinct lines (10°C, 15°C, 20°C, 25°C, and 30°C), illustrates unique trends in nymph production. At the lowest temperature of 10°C, nymph production starts at about 1 nymph per female per day, fluctuates, and peaks around day 3 before gradually declining. This temperature maintains relatively consistent, albeit declining, reproductive output until it drops sharply after day 20. The reproductive output at 15°C shows initial fluctuations with several peaks and valleys but generally trends higher compared to 10°C, it peaks (average 5 nymphs/female/day) around day 19, after which it drops significantly. At 20°C, the pattern starts similarly with fluctuations, reaching several peaks which are higher than those observed at cooler temperatures. The reproductive output remains relatively high and stable until it begins a steep decline around day 18. In the 25°C condition, nymph production starts higher than in all cooler conditions, peaks early, and maintains a high output longer, until a sharp decline after day 15, suggesting that while higher temperatures may initially promote greater reproductive output, they also lead to quicker declines. At the highest temperature of 30°C, nymph production is initially low, peaks slightly around day 7, and falls to zero by day 12, indicating that extremely high temperatures may severely inhibit reproductive capacity.

### e) Life table parameters of *Myzus persicae*

Table 2 provides detailed life table parameters of the aphid *M. persicae* across a range of temperatures from 10°C to 30°C, illustrating the effects of temperature on reproductive and survival metrics. At 10°C, the net reproduction rate (R_o_) is relatively low at 170.36, and it increases significantly at 15°C to 365.80, indicating more favourable conditions for reproduction at this slightly warmer temperature. However, as the temperature increases further to 20°C and 25°C, R_o_ showed varied responses, with a decrease at 20°C (219.48) and a moderate increase at 25°C (272.30), followed by a drastic drop at 30°C (51.30). The intrinsic rate of increase (r_m_) also varied with temperature, starting at 0.76 at 10°C and peaking at 0.98 at 30°C. The finite rate of increase (λ) follows a similar trend, gradually increasing with temperature. Mean generation time (T) decreased as temperature increases, from 6.84 days at 10°C to 3.96 days at 30°C, showing that higher temperatures accelerate life cycle completion. Doubling time (DT), which indicates the time it takes for the population to double, also decreases with rising temperature, from 0.93 days at 10°C to 0.71 days at 30°C.

**Table 2.** Mean ± SE of life table parameters of aphid *Myzus percicae* under different temperatures.

| Temperature (°C) | Net reproduction rate ($R_0$) | Intrinsic rate of increase ($r_m$) | Finite rate of increase ($\lambda$) | Mean generation time ($T$) | Doubling time (DT) |
| --- | --- | --- | --- | --- | --- |
| 10°C | 170.36 ± 16.4 **a** | 0.76 ± 0.03 **a** | 2.14 ± 0.07 **a** | 6.84 ± 0.3 **ab** | 0.93 ± 0.04 **a** |
| 15°C | 365.80 ± 37.1 **b** | 0.77 ± 0.02 **a** | 2.16 ± 0.05 **a** | 7.69 ± 0.3 **a** | 0.91 ± 0.03 **a** |
| 20°C | 219.48 ± 35.9 **ac** | 0.96 ± 0.04 **ab** | 2.62 ± 0.12 **b** | 5.66 ± 0.3 **c** | 0.74 ± 0.03 **b** |
| 25°C | 272.30 ± 37.8 **abc** | 0.85 ± 0.03 **bc** | 2.36 ± 0.07 **ab** | 6.56 ± 0.3 **bc** | 0.82 ± 0.03 **ab** |
| 30°C | 51.30 ± 4.7 **c** | 0.98 ± 0.05 **c** | 2.78 ± 0.15 **b** | 3.96 ± 0.2 **d** | 0.71 ± 0.04 **b** |
| | $\chi^2 = 32.47$ | F = 8.44 | $\chi^2 = 21.26$ | F = 24.24 | F = 8.63 |
|  | df = 4 | df = 4 | df = 4 | df = 4 | df = 4 |
|  | p <0.0001 | p <0.0001 | p <0.0001 | p <0.0001 | p <0.0001 |

### f) *Potato virus Y* acquisition and transmission by *Myzus persicae* to potato plants

The temperature-dependent transmission rate of PVY by the aphid *M. persicae* was best captured by the Taylor model, which predicted the highest transmission rate between 15°C and 20°C (Figure 5A and Table S6). This optimal range of temperatures yields the highest transmission rate. At temperatures below and above this range, the transmission rate declines, with the lowest rates observed near 0°C and 45°C. Figure 5B presents the interaction between the transmission rate of Potato virus Y (PVY), survival, and total oviposition of *M. persicae* across different temperatures. The transmission rate of PVY increases with temperature, peaking around 18°C, and declines at higher temperatures, indicating optimal transmission efficiency at moderate temperatures. The survival rate of *M. persicae* reaches its maximum between 20°C and 25°C, decreasing significantly at both lower and higher temperatures. Similarly, fecundity, or total oviposition, peaks between 18°C and 22°C and diminishes outside this optimal range, showing that reproductive output is highest at moderate temperatures. Notably, the optimal temperature for the highest PVY transmission rate slightly precedes the peak survival and fecundity rates of *M. persicae*. This suggests that while moderate temperatures favour virus transmission, slightly warmer temperatures are more conducive to aphid survival and reproduction.

**Figure 5.**
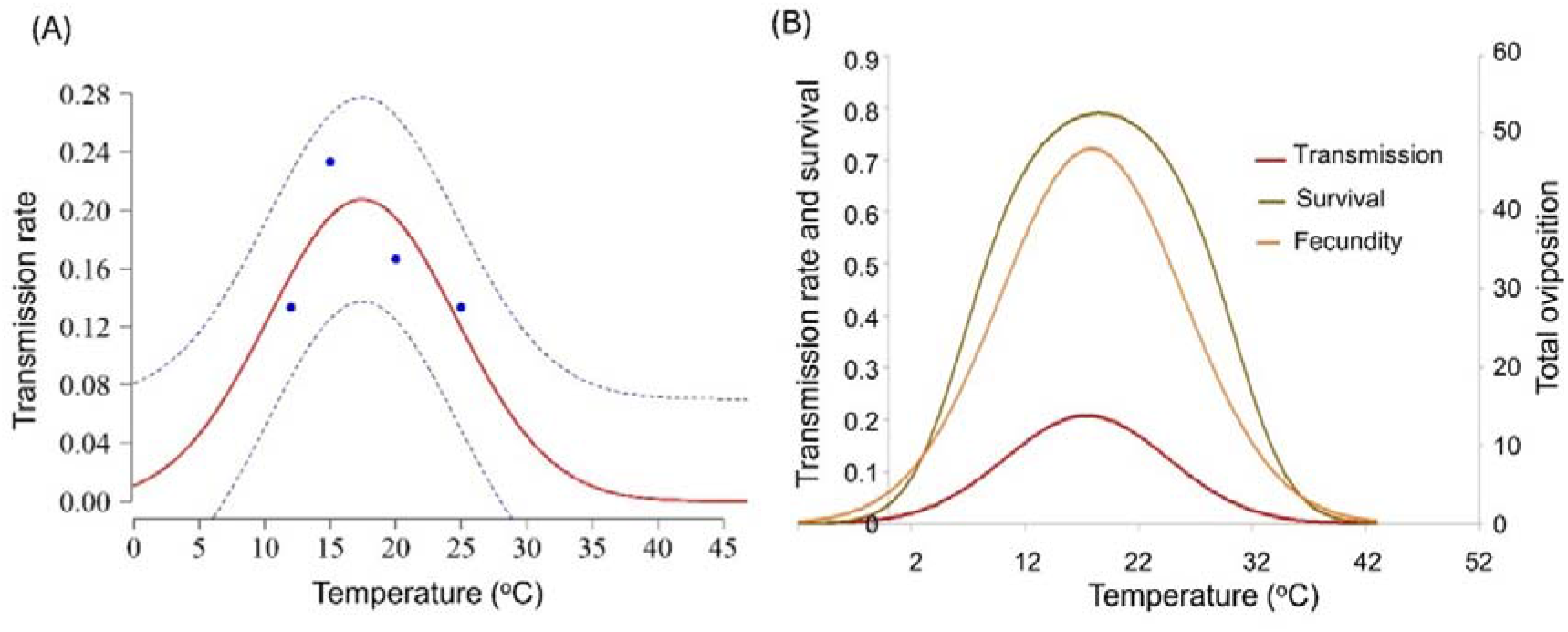
Transmission rate of Potato virus Y (PVY) (A) and its interaction with survival and total oviposition (B) of aphid *M. persicae*

### g) Suitable Establishment Risk Index (ERI), Generation Risk Index (GI) and potential Potato virus Y transmission across the globe

Figure 6 illustrates the Establishment Risk Index (ERI) (A) and Generation Risk Index (GRI) (B) of *M. persicae* in potato growing regions based on global climate temperature data. Regions with the highest ERI (0.90-1.00) indicate a very high likelihood of *M. persicae* establishment. These areas include tropical and subtropical regions across South America, Africa, South Asia, Southeast Asia, and parts of Oceania. Moderate-risk regions (0.40-0.80) are found in temperate zones, including parts of North America, Europe, and East Asia. Low-risk areas (0.10-0.30) are primarily located in higher latitudes and mountainous regions with cooler temperatures. Very low-risk zones (0.00-0.10) are found in extremely cold climates such as the Arctic, Antarctic, and high-altitude regions (Figure 6B).

**Figure 6.**
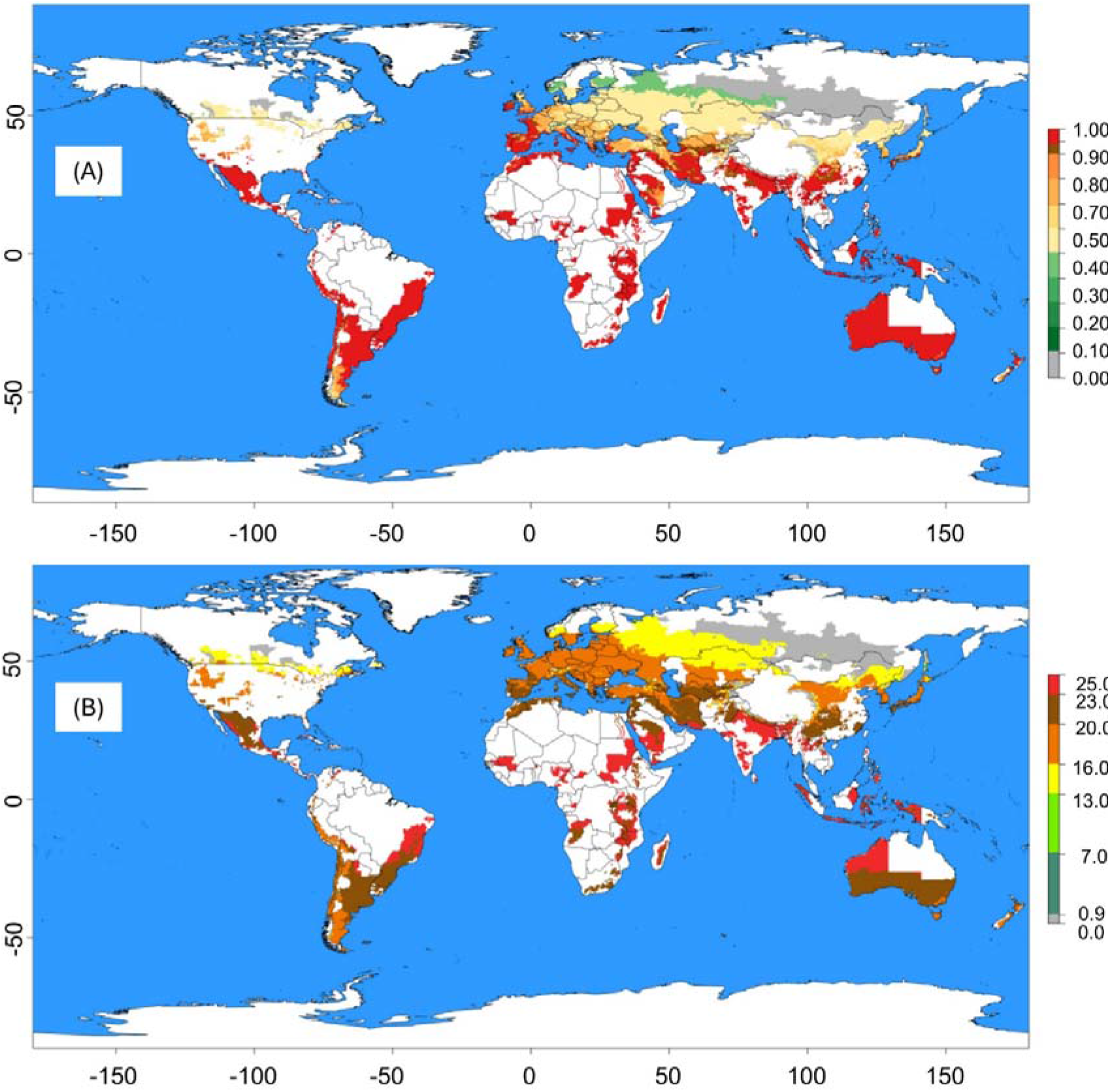
Establishment Risk Index (A) and Generation Risk Index (B) of *M. persicae* prediction for the whole world according to climate temperature data from WorldClim

Regions with the highest GI (23.0-25.0 generations) indicate a high potential for multiple generations per year. These areas are predominantly in tropical and subtropical regions of South America, Africa, South Asia, Southeast Asia, and parts of Oceania. Moderate GI regions (13.0-20.0 generations) are mainly in temperate zones, including parts of North America, Europe, and East Asia. Low GI areas (7.0-13.0 generations) are in higher latitudes and mountainous regions with cooler temperatures. Very low GI zones (0.0-7.0 generations) are found in extremely cold climates like the Arctic, Antarctic, and high-altitude regions (Figure 6B).

Figure S2 illustrates the Potential Activity of Transmission (PAT) of Potato virus Y by *M. persicae* based on global climate temperature data. Regions with the highest PAT values (1.2-1.4) indicate a very high potential for virus transmission, primarily located in tropical and subtropical regions of South America, Central Africa, South Asia, and Southeast Asia. Areas with moderate PAT values (0.6-1.2) are found in parts of North America, South America, Africa, Europe, and East Asia, exhibiting a significant potential for virus transmission. Regions with low PAT values (0.2-0.6) are mainly situated in temperate zones, including parts of North America, Europe, and East Asia, while very low PAT areas (0.0-0.2) are typically located in extremely cold climates, such as the Arctic, Antarctic, and high-altitude regions (Figure S2).

## 4. Discussion

Although many unknows remain about specific effects of global warming, it is widely accepted that it significantly affects crops, such as potatoes, as well as the insect pests associated with them, like aphids. This is likely through the effects of climatic variables, especially temperature. Temperature is a critical factor influencing the lifespan, survival, and development of the green peach aphid, *Myzus persicae*. Our study demonstrates that the developmental time of *M. persicae*, including nymph and adult longevity, is significantly affected by temperature. The longest life span was observed at the lowest temperatures (10°C and 15°C), while the shortest lifespan occurred at the highest temperature (30°C). The study by Baral et al., (2022), which assessed the effect of temperatures ranging from 17°C to 29°C on *M. persicae*, also found that longevity decreases as temperature increases. However, their study reported longer lifespans for *M. persicae* compared to ours, which may be attributed to their use of cabbage (*Brassicae olerasia*) as a host plant [11]. Additionally, the study of Khurshid et al. (2022) showed that higher temperatures are detrimental to the survival and reproduction of *M. persicae* [10].

The survival and life expectancy trends across different temperatures reveal that cooler temperatures favour higher survival rates and longer life expectancy for both nymph and adult stages. At 10°C, nymphs exhibited nearly 100% survival up to around day 9, followed by a gradual decline until reaching 0% by day 19. In contrast, survival at 30°C plummeted to 0% by day 15 (Figure 2A). This indicates that extreme temperatures, particularly high temperatures, have a detrimental impact on aphid survival. Multiple studies have concluded that temperature extremes affect insect development, fecundity, survival, and geographic distribution [7,29,30]. At 10°C, adult aphids maintain over 50% survival up to about day 12, with a gradual decline to approximately 20% by day 17. As temperatures increase, the survival rate declines more rapidly, dropping to below 10% by day 9 at 30°C (Figure 3B). These findings suggest that while moderate temperatures favour aphid longevity, higher temperatures significantly reduce their life expectancy. This underscores the need for temperature-specific management strategies in pest control [9].

Fecundity of *M. persicae* demonstrates a classic hump-shaped relationship with temperature. At 10°C, the average fecundity is 29.81 nymphs per female, with a peak at 15°C (45.82 nymphs/female). After this point, fecundity decreases, with the lowest value (14.25 nymphs per female) at 30°C. This pattern suggests that intermediate temperatures favour higher reproductive output. The relationship between temperature and fecundity was best described by the Wang 11 model, which predicted the highest fecundity at 15-20°C. Life table parameters of *M. persicae*, including net reproduction rate, intrinsic rate of increase, and mean generation time, are significantly influenced by temperature. The net reproduction rate is highest at 15°C, indicating favourable conditions for reproduction at this temperature. However, as temperature increases beyond this point, net reproduction rate decreases, with the lowest value observed at 30°C. The intrinsic rate of increase peaks at 30°C, suggesting that the population growth rate is highest at the warmest temperature despite the lower net reproduction rate. These findings highlight the complex interactions between temperature and aphid population dynamics, underscoring the need for predictive models to inform pest management practices under varying climatic conditions [13,30].

These observations are consistent with previous studies that have documented the impact of temperature on aphid reproduction. For instance, the study by Ahn et al. (2020) on *Acyrthosiphon pisum* (pea aphid) showed highest fecundity at 15°C and 20°C (74.9 and 62.5 nymphs/female, respectively) and significantly declining as temperature increases with the lowest at 30°C (4.5 nymphs/female) using *Vicia faba* as a host plant [31]. The study of Cividanes and Souza (2003) in *M. persicae* showed highest fecundity at 15°C (69.2 nymphs/female) and lowest fecundity at 25°C (30.7 nymphs/female) using *B. oleracea* as a host plant [32]. However, the study of Baral et al. (2022) showed higher fecundity of *M. persicae* at 23°C (72.4 nymphs/female); fecundity decreased below and upper this threshold with the lowest at 29°C (28 nymphs/female) in *B. oleracea* [11]. These findings are further supported by the work of Leather et al. (1989), who reported that temperature and host specie play a critical role in the reproductive success of aphids. Their research found that the aphid *Rhopalosiphum padi* fecundity is higher at 15°C when the host are either oats cv. Aster (38.2 nymphs/female) or timothy grass (33.6 nymphs/female) and fecundity decreases when temperature is 20°C when the host is either oats cv. Aster (28.47 nymphs/female) or wheat cv. Huntsman (25.14 nymphs/female) [33]. Moreover, other studies have explored the physiological mechanisms underlying this temperature-dependent fecundity pattern. Dixon (1977) suggested that lower temperatures might prolong the developmental period of aphids, allowing them more time to produce offspring [34]. Conversely, higher temperatures may accelerate development but at the cost of reduced fecundity and longevity.

Our study reveals that the efficiency of virus transmission is closely linked to the aphids’ physiological state, which is temperature dependent (Figure 5B). Ng & Perry (2004) demonstrated that moderate temperatures enhance aphid feeding activity and mobility, thereby increasing the likelihood of virus transmission [35]. We found that PVY transmission by *M. persicae* was highest at 20°C, supporting the optimal physiology and behaviour for virus transmission. In comparison, Nancarrow et al. (2014) showed that the transmission of Barley yellow dwarf virus (BYDV), which is persistently transmitted by *Rhopalosiphum padi*, was highest at temperatures 10-21.1°C (night-day fluctuations) compared to 5-16°C (night-day fluctuation) [36]. The temperature-dependent pattern of PVY transmission by *M. persicae* can likely be attributed to physiological and behavioural mechanisms. At optimal temperatures, aphids exhibit increased feeding frequency and duration, enhancing virus acquisition and transmission. van Emden & Harrington (2017) explained that temperature affects the metabolic rate of aphids, influencing their feeding behaviour and transmission efficiency [37]. Additionally, (van Munster, 2020) noted that the stability and infectivity of plant viruses within aphid vectors can be temperature-dependent, with moderate temperatures maintaining virus viability and extreme temperatures degrading viral particles, reducing transmission efficiency [38]. Besides that, PVY is transmitted by aphids through direct reversable binding of virus particles to the aphid stylet during feeding and salivation with the help of the virus encoded HC-Pro protein [39] and thus temperature may also affect the stoichiometry of this binding process to influence transmission efficiency. Although in this study temperature dependent transmission efficiency closely resembled those of aphid life table parameters, in a study of potato yellow vein virus (PYVV) transmission by the greenhouse whitefly, transmission efficiency followed a dissimilar trend than the whiteflies life parameters [28], indicating this can vary among virus-vector system and may need to be studied individually for each case.

Using the risk prediction tool from the ILCYM 4.0 software, we mapped the potential activity of PVY transmission by *M. persicae* based on global climate temperature data. Regions with the highest potential activity are primarily located in tropical and subtropical regions, including South America, Central Africa, South Asia, and Southeast Asia. Tropical and subtropical regions, which exhibit higher average temperatures and humidity, provide ideal conditions for *M. persicae* activity and PVY transmission. Studies by Radcliffe & Ragsdale (2002) and Ragsdale et al. (2001) confirm that warmer climates enhance aphid population growth and virus transmission rates [2,40]. The presence of diverse and abundant host plants in these regions further supports the establishment and proliferation of both the aphid vector and the virus. Similarly, Gamarra et al. (2020) highlighted the increased risk of PVY transmission in tropical regions due to the favourable climate for both the whitefly vector and virus persistence [28]. Moderate-risk regions include parts of North America, Europe, and East Asia. These temperate zones experience seasonal variations that can influence aphid population dynamics and virus transmission. In these regions, warmer months provide suitable conditions for *M. persicae* activity, while cooler periods may limit their population growth and transmission potential. Bhoi et al. (2022) indicated that in temperate climates, integrated pest management strategies are essential during peak aphid activity periods to mitigate the risk of PVY spread [41]. Low-risk areas are situated in temperate zones and extremely cold climates such as the Arctic and Antarctic. These regions have limited aphid activity due to unfavourable temperatures for *M. persicae* survival and reproduction. Extreme cold climates inhibit aphid population growth and virus transmission [33,34]. The short growing seasons and harsh environmental conditions in these regions restrict the establishment and spread of PVY.

The temperature-dependent fecundity pattern of *M. persicae* has significant implications for integrated pest management (IPM) strategies. Understanding the optimal and suboptimal temperature ranges for aphid reproduction aids in predicting population dynamics and devising effective control measures. During moderate temperature periods, aphid populations may surge, necessitating timely interventions like biological control agents or selective insecticides, while extreme temperatures may naturally limit reproduction, reducing the need for intensive pest control. Our study emphasizes the importance of temperature-specific management strategies, incorporating certified virus-free seed potatoes, crop rotation, insecticides, and PVY-resistant potato varieties [42,43]. Biological control agents, such as natural predators and parasitoids, should also be integrated into IPM programs. Additionally, understanding geographical variations in PVY transmission risk is critical for effective IPM strategies. In high-risk regions, continuous monitoring and early detection are essential, while in moderate-risk areas, targeted interventions during peak aphid activity can manage virus transmission. Implementing predictive models that incorporate climate data can forecast high-risk periods and guide control measures.

## Conclusion

Our study reveals that temperature significantly affects the life history traits and PVY transmission efficiency of *M. persicae*, with optimal development and highest fecundity at lower temperatures and peak virus transmission at 20°C. Using the ILCYM software, we identified high-risk PVY transmission regions in tropical and subtropical areas, moderate-risk in temperate zones, and low-risk in extremely cold climates. These temperature-dependent patterns underscore the need for integrated pest management (IPM) strategies tailored to specific temperature ranges. Effective control measures include the use of certified virus-free seed potatoes, crop rotation, insecticides, PVY-resistant varieties, and biological control agents. The study highlights the importance of predictive models that incorporate ecological and epidemiological data to inform pest management under changing climatic conditions, aiming to mitigate the impact of *M. persicae* and PVY on potato production, thereby supporting sustainable agriculture and global food security.

## Acknowledgments

This study was supported by the Earth Commons, Georgetown University’s Institute for Environmental and Sustainability. The authors thank the Eco Impact Awards for providing the funding to carried out these experiments. The authors also want to thank the International Centre for Insect Physiology and Ecology (*icipe*) and the International Potato Center (CIP) for facilitating their laboratories and personnel to perform the above-mentioned experiments. The authors at *icipe* gratefully acknowledge *icipe* core financial support organizations and agencies: the Swedish International Development Cooperation Agency (Sida); the Swiss Agency for Development and Cooperation (SDC); the Australian Centre for International Agricultural Research (ACIAR); the Norwegian Agency for Development Cooperation (Norad); the German Federal Ministry for Economic Cooperation and Development (BMZ); and the Government of the Republic of Kenya.

## Funding

This research was funded by the Eco Impact Awards to OCV, AA, and PAA. HG, PC, and JK were supported by the CGIAR initiative on plant health supported by the donors of the CGIAR Trust Fund (http://www.cgiar.org/funders/). LRJ was partially supported by NSF DMS/DEB grant #1750113.

## Availability of data and materials

R code for creating model outputs and data generated for adult survival, nymph survival, longevity, fecundity, nymph life expectancy, adult life expectancy for *Myzus persicae* (the green peach aphid) are publicly available on Zenodo at https://zenodo.org/records/13858882. Supplementary material is available online.

## Notes

### Competing Interest Statement

The authors have declared no competing interest.

