## Supplementary material for "Effects of temperature on the life-history traits of *Myzus persicae* and its efficiency in transmitting potato virus Y (PVY) in potato crops": S1

- Corresponding author

**Table S1:** Nymphal and adult development time and mortality rate for *Myzus persicae* under different temperatures using the Insect Life Cycle Modeling (ILCYM) software.

| Temp. |  |  | Nymphs | |  |  | Adult | |
| --- | --- | --- | --- | --- | --- | --- | --- | --- |
|  | N ^A^ |  | Median dev. time | | Mortality |  | Median dev. time | |
| (°C) |  |  | (days) ^C^ | | (%) |  | (days) | |
| 10 | 100 |  | 12.83 (0.824) | | 42 |  | 11.43 (0.872) | |
| 15 | 100 |  | 11.27 (0.944) | | 20 |  | 11.69 (1.132) | |
| 20 | 100 |  | 10.11 (0.902) | | 32.2 |  | 9.59 (0.977) | |
| 25 | 100 |  | 10.36 (0.874) | | 22 |  | 10.69 (1.065) | |
| 30 | 100 |  | 11.47 (1.182) | | 64 |  | 6.06 (0.711) | |
|  | Model ^B^ |  | Weibull | |  |  | Weibull | |
|  | *ln*(scale) |  | -2.959 (0.043)*** | |  |  | -0.9738 (0.0470)*** | |
|  | Scale δ |  | 0.0519 (0.002)*** | |  |  | 0.378 (0.018)*** | |
|  | α = 1/ δ |  | 19.3 (0.84)*** | |  |  | 2.64 (0.124)*** | |
|  | |  | *ln* L | ΔDeviance |  |  | *ln* L | ΔDeviance |
| Intercept only | |  | -757.6 |  |  |  | -871.6 |  |
| λ for each Temp. | |  | -731.7 | 1469.38 |  |  | -842.9 | 142.32 |
| P | |  | <0.001 | | 0.2472 |  | <0.001 | |

^A^	N is the number of individuals evaluated at a given temperature.

^B^	δ is the scale of the selected distribution link function; the figures in () are SE of *ln*(δ), δ, and α (“***” indicates P < 0.001). The accumulated development frequency in relation to normalized age (time/median time) is calculated according to the selected distribution link function; for example, for the log-logistic link function: accu. dev. freq. = 1-(1/(1+*x^α^*)), where *x* is the normalized age (determined through rate summation), and *α* = 1/δ.

^C^	Numbers in parenthesis are 95% confidence limits based on t-distribution (a heterogeneity factor, H = deviance/df, was included to calculate the limits). Medians followed by different letters in the same columns are significantly different (P < 0.05) according to the AFT mode

**Table S2:** Models and their parameters fitted to describe the development rate (1 per day) for immature life stages of *M. persicae* reared on potato plants.

| Life Stages | Parameter estimates of the model^A^ | | | *F* value | *df* _1,2_ | *P* | AICc |
| --- | --- | --- | --- | --- | --- | --- | --- |
|  | *T_opt_* | *T_roh_* | *r_m_* |  |  |  |  |
| Nymph | 25.75 (±1.194)** | 2.66 (±0.644)** | 0.004 (±0.002)* | 86.37 | 2, 2 | 0.011 | -17.107 |

Numbers in parenthesis are standard errors. Parameter values significantly different from zero are indicated by asterisks (P < 0.05 = *, P < 0.01 = **, P < 0.001 = ***).^A^ The equation of the Taylor model is:

$$r(T)=r_{m}*e^{-\frac{1}{2}\left( -\frac{\left( T-T_{opt} \right)}{T_{roh}} \right)^{2}}$$

where *r*(*T*) is the development rate at temperature *T*, then *T_opt_* , *T_roh_* and *r_m_*, are parameters of the equation.

**Table S3:** Models and their parameters fitted to describe the mortality rate for immature life stages of *M. persicae* reared on potato plants.

| Life stages | Parameter estimates of the Quadratic model^A^ | | | *F*-value | *df*_1, 2_ | *P* | AICc |
| --- | --- | --- | --- | --- | --- | --- | --- |
|  | *a* | *b* | *c* |  |  |  |  |
| Nymph | 0.013 (±0.71)** | -0.492 (±0.242)** | 3.25 (±2.225)* | 3.04 | 2, 2 | 0.247 | 11.8 |

Numbers in parenthesis are standard errors. Parameter values significantly different from zero are indicated by asterisks (P < 0.05 = *, P < 0.01 = **, P < 0.001 = ***). **^A^** The equation of the Quadratic model is:

$$\boldsymbol{m}\left( \boldsymbol{T} \right)\boldsymbol{=a+b(T)+ c(}\boldsymbol{T}^{\boldsymbol{2}}\boldsymbol{)}$$

where *m*(*T*) is the mortality at temperature *T*, then *a*, b and *c*, are parameters of the equation.

**Table S4:** Median oviposition time and mean fecundity for *Myzus persicae* under different temperatures

| Temp. |  |  | Median oviposition time | |  | Mean fecundity |
| --- | --- | --- | --- | --- | --- | --- |
| (°C) |  |  | (days)(±SE) | |  | (eggs/female) (±SE) |
| 10 | 50 |  | 5.87 (0.839) | |  | 30.16 (1.919) |
| 15 | 79 |  | 6.51 (1.116) | |  | 46.25 (2.405) |
| 20 | 62 |  | 4.99 (0.921) | |  | 39.68 (2.084) |
| 25 | 69 |  | 5.74 (1.036) | |  | 39.52 (2.598) |
| 30 | 36 |  | 3.09 (0.916) | |  | 14.66 (1.107) |
|  | Model ^B^ |  | Weibull | |  |  |
|  | *ln*(scale) |  | -0.4391 (0.007)*** | |  |  |
|  | Scale δ |  | 0.6446 (0.004)*** | |  |  |
|  | α = 1/ δ |  | 1.5513 (0.012)*** | |  |  |
|  | |  | *ln* L | ΔDeviance |  |  |
| Intercept only | |  | -34221.6 |  |  |  |
| λ for each Temp. | |  | -33927.7 | 7182.3 |  |  |
| P | |  | <0.001 | |  | 0.079 |

^A^	N is the number of individuals evaluated at a given temperature.

^B^	δ is the scale of the selected distribution link function; the figures in () are SE of *ln*(δ), δ, and α (“***” indicates P < 0.001). The accumulated development frequency in relation to normalized age (time/median time) is calculated according to the selected distribution link function; for example, for the log-logistic link function: accu. dev. freq. = 1-(1/(1+*x^α^*)), where *x* is the normalized age (determined through rate summation), and *α* = 1/δ.

**Table S5:** Models and their parameters fitted to describe adult senescence rate, total number of eggs per female, and oviposition time for *M. persicae* reared on potato plants.

| Response variable | Models ^a^ | Parameters | *F* value | df _1,2_ | *P* | AICc |
| --- | --- | --- | --- | --- | --- | --- |
|  |  | $b_{0}$ 0.139 (±0.002) ^b^ | 3.523 | 2,2 | 0.221 | 30.927 |
| Adult senescence rate | Quadratic *s*$\left( \boldsymbol{T} \right)\boldsymbol{=}b_{0}+b_{1}(T)+ b_{2}( \boldsymbol{T}^{\boldsymbol{2}}\boldsymbol{)}$ | $b_{1}$ -0.007 (±0.007) |  |  |  |  |
|  |  | $b_{2}$ 0.0003 (±0.0002) |  |  |  |  |
| Total eggs per female | Quadratic *t*$\left( T \right)=b_{0}+b_{1}(T)+ b_{2}( \boldsymbol{T}^{\boldsymbol{2}}\boldsymbol{)}$ | $b_{0}$ -0.0076 (±0.0705) | 11.639 | 2,2 | 0.0791 | 0.5821 |
|  |  | $b_{1}$ 0.2754 (±0.078) |  |  |  |  |
|  |  | $b_{2}$ 1.4069 (±0.724) |  |  |  |  |
| Oviposition time^-1^ | Taylor $o(T)=r_{m}*e^{-\frac{1}{2}\left( -\frac{\left( T-T_{opt} \right)}{T_{roh}} \right)^{2}}$ | $T_{opt}$ 15.109 (±3.762) | 3.436 | 2,2 | 0.225 | 24.028 |
|  |  | $T_{roh}$ 16.407 (±5.93) |  |  |  |  |
|  |  | $r_{m}$ 1.8456 (±0.127) |  |  |  |  |

a Models: Quadratic: s(T) is the senescence rate at temperature T (°C), and b is equation parameters. Quadratic: t(T) represents the fecundity function at temperature T (°C) and b_1_, b_2_, and b_3_ are parameters of the equation, and o(T) is the inverse oviposition time where T_opt_, T_roh_, and r_m_ are parameters of the equation.

b Numbers in parenthesis are standard errors.

**Table S6:** Models and their parameters fitted to describe the acquisition of potato virus Y (PVY) by *Myzus persicae* in potato plants

| Response variable | Models ^a^ | Parameters | *F* value | df _1,2_ | *P* | AICc |
| --- | --- | --- | --- | --- | --- | --- |
| Adult senescence rate |  | $T_{opt}$ 17.435 (±0.688)^b^ | 5.871 | 2,9 | 0.023 | -5.182 |
|  | Taylor: $m\left( T \right)=1-rm*e^{\left( -\frac{1}{2}\left( -\frac{\left( T-T_{opt} \right)}{T_{roh}} \right)^{2} \right)}$ | $T_{roh}$ 7.177 (±1.172) |  |  |  |  |
|  |  | $r_{m}$ 0.2077 (±0.016) |  |  |  |  |

a Models: Taylor: m(T) is the transmission rate function, where T_opt_, T_roh_, and r_m_ are parameters of the equation.

b Numbers in parenthesis are standard errors.

**Figure S1:** *Myzus persicae* Potential Activity of Transmission (PAT) of Potato virus Y according to climate temperature data from WorldClim.

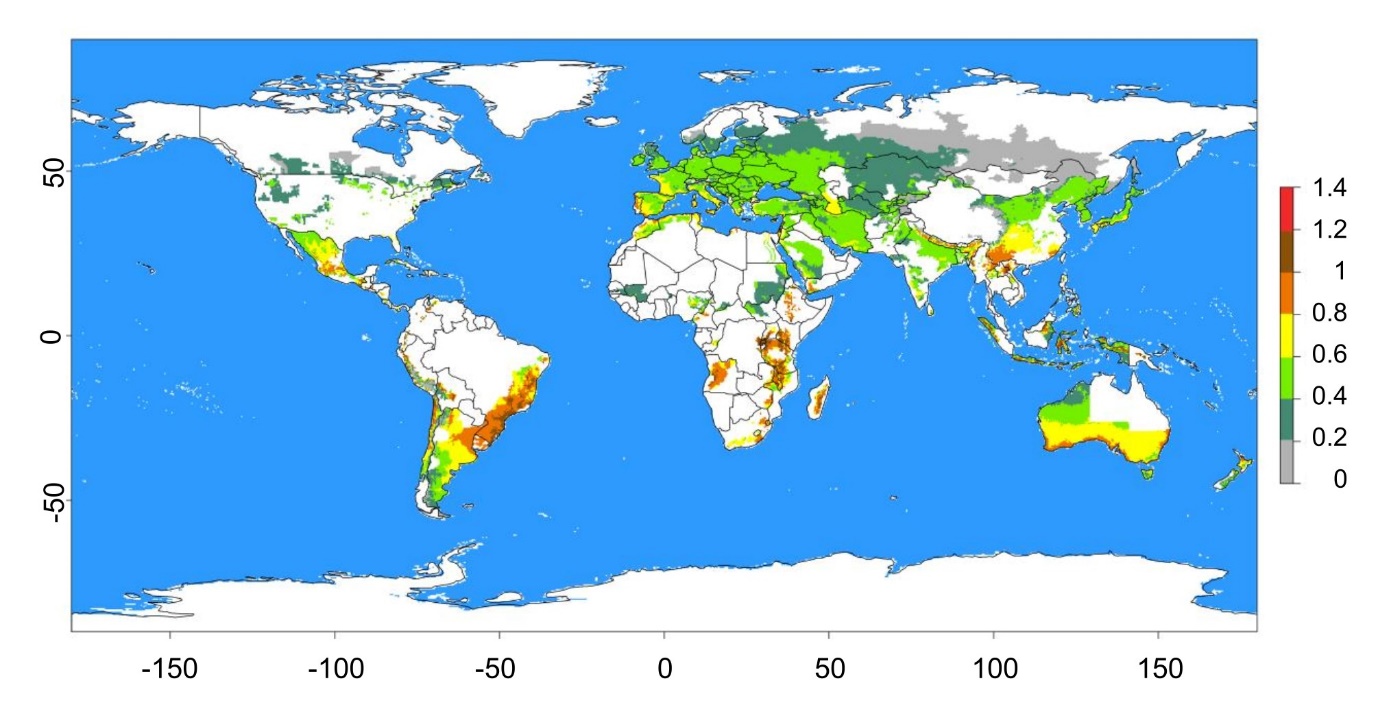

**Figure S2:** Diagram showing the procedure of virus transmission experiments of potato virus Y (PVY) by Myzus persicae under constant temperatures (12°C,15°C, 20°C, 25°C).

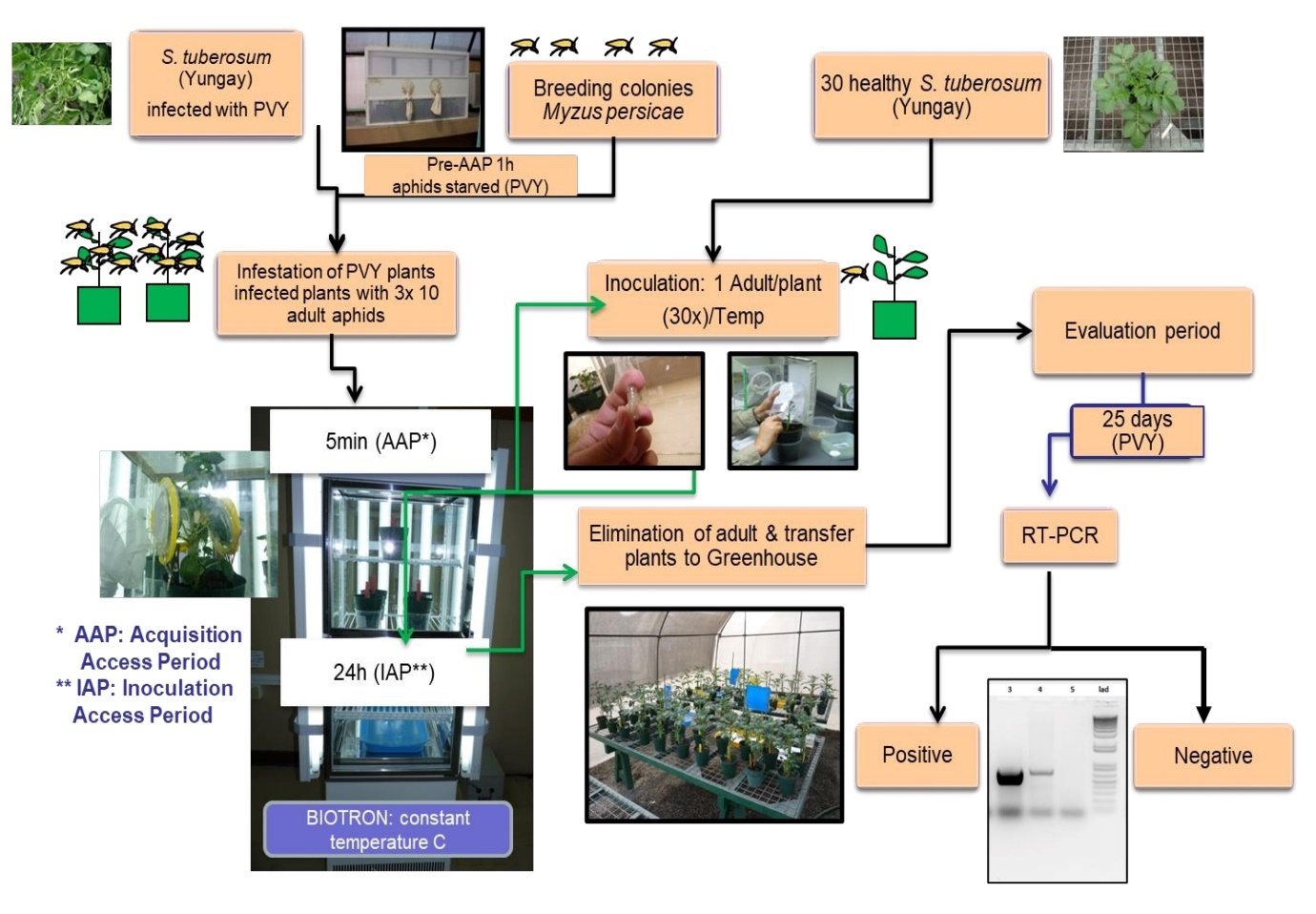
